# Primed and ready: Nanopore metabarcoding can now recover highly accurate consensus barcodes that are generally indel-free

**DOI:** 10.1101/2023.08.04.552069

**Authors:** Jia Jin Marc Chang, Yin Cheong Aden Ip, Wan Lin Neo, Maxine A. D. Mowe, Zeehan Jaafar, Danwei Huang

**Affiliations:** Department of Biological Sciences, National University of Singapore, 16 Science Drive 4, Singapore 117558, Singapore; Animal & Veterinary Service, National Parks Board, 1 Cluny Road, Singapore 259569; Lee Kong Chian Natural History Museum, National University of Singapore, 2 Conservatory Drive, Singapore 117377, Singapore; Tropical Marine Science Institute, National University of Singapore, 18 Kent Ridge Road, Singapore 119227, Singapore; Centre for Nature-based Climate Solutions, National University of Singapore, 6 Science Drive 2, Singapore 117546, Singapore

**Keywords:** Biomonitoring, Illumina, MinION, Next-generation sequencing, Species diversity, Zooplankton

## Abstract

**Background:** DNA metabarcoding applies high-throughput sequencing approaches to generate numerous DNA barcodes from mixed sample pools for mass species identification and community characterisation. To date, however, most metabarcoding studies employ second-generation sequencing platforms like Illumina, which are limited by short read lengths and longer turnaround times. While third-generation platforms such as the MinION (Oxford Nanopore Technologies) can sequence longer reads and even in real-time, application of these platforms for metabarcoding has remained scarce due to the relatively high read error rate as well as the paucity of specialised software for processing such reads.

**Findings:** We show that this is no longer the case by performing nanopore-based metabarcoding on 34 zooplankton bulk samples with amplicon_sorter, benchmarking the results against conventional Illumina MiSeq sequencing. The R10.3 sequencing chemistry and super accurate (SUP) basecalling model reduced raw read error rates to ∼4%, and consensus calling with amplicon_sorter (without further error correction) generated metabarcodes that were ≤1% erroneous. Although Illumina recovered a higher number of molecular operational taxonomic units (MOTUs) than nanopore sequencing (589 vs. 471), we found no significant differences in the zooplankton communities inferred between the sequencing platforms. Indeed, the same ecological conclusions were obtained regardless of the sequencing platform used. Moreover, 406 of 444 (91.4%) shared MOTUs between Illumina and nanopore were found to be indel-free.

**Conclusions:** Collectively, our results illustrate the viability of nanopore metabarcoding for characterising communities, and paves the way for greater utilisation of nanopore sequencing in various metabarcoding applications.

## Background

DNA metabarcoding refers to the high-throughput sequencing of total (and sometimes degraded) DNA from bulk or environmental samples (e.g., air, water, soil, faeces, etc.) with the goal of multispecies identification [1]. It was built upon the DNA barcoding paradigm that has been established for about two decades involving the sequencing of short segments of DNA (termed “barcodes”) and matching them to sequence databases to obtain species identities [2]. DNA metabarcoding emerged in the 2010s, and was primarily made possible due to rapid advancements in nucleic acid sequencing technologies—with “next-generation sequencing” (NGS) platforms—which have the ability to generate billions of sequence reads in a single experiment [3]. This development has been groundbreaking due to the sheer ability of NGS platforms to generate sequence reads (i.e., DNA barcodes) in parallel, so multispecies detections and identification from various sample types are now possible. This has led to a meteoric rise in the number of studies that have since performed NGS-based barcoding or metabarcoding for various applications. For instance, 60% of DNA sequencing studies in marine science published yearly between 2013 and mid-2022 generated their sequence reads with Illumina [4]. The release of the MinION in 2014 by Oxford Nanopore Technologies (ONT) became another significant milestone in nucleic acid sequencing for several reasons: (1) its lower entry and per-base sequencing cost (1,000 USD for the entry starter pack), (2) its ability to perform long-read sequencing (now up to ∼4 Mb long), (3) its compact size and portability, and (4) its ability to generate data in real-time [5,6]. All these were perhaps a direct response to common criticisms of Illumina sequencing, which was comparatively more expensive, and limited by its short read-lengths (up to ∼500 bp). Since then, nanopore sequencing has been applied in numerous whole-genome sequencing studies [7–10] and metagenomic studies [11,12].

However, nanopore metabarcoding applications remain relatively uncommon, and this is evident in the low number of published papers, especially in biodiversity-related fields. Such studies focused on microbes [13–17], and few have paid attention to non-microbial taxa until more recently. Importantly, Krehenwinkel et al. [18] and Baloğlu et al. [19] laid the groundwork with ONT’s MinION sequencer by successfully metabarcoding mock communities comprising nine arthropod and 50 aquatic invertebrate species respectively. Other studies have since applied nanopore metabarcoding for biodiversity and community characterisation [14,20–23], species-specific detections [24,25], and even diet analysis [26] with actual samples. The consensus from the abovementioned studies is that nanopore sequencing shows promise in metabarcoding.

We posit that the general lack of nanopore-based metabarcoding studies can be attributed to two main factors. The first is the relatively high error rate in nanopore reads; early studies have reported error rates of ∼20% [27] to as high as 38% [28]. In contrast, the error rate of Illumina sequencing is only 0.24% [29]. There is thus concern that the high error rates would hinder accurate species identification in DNA metabarcoding. The second factor could be the lack of programs to process nanopore reads for metabarcoding (but see below), compared to the plethora of pipelines catered to short-read sequencing, like APSCALE [30], DADA2 [31], eDNAflow [32], or OBITools [33]. DADA2 currently supports PacBio circular consensus sequencing but not nanopore reads [34], and even ONT’s EPI2ME platform is intended for microbial sequencing only. Nanopore-specific workflows like ONTrack [35], NGSpeciesID [36] and miniBarcoder [37,38] were designed mainly for DNA barcoding, although Davidov et al. [14] have successfully applied ONTrack to process their metabarcoding reads. Prior metabarcoding studies have worked around the lack of specialised software by either (i) conducting BLAST searches of raw nanopore reads with stricter e-value settings as low as 1e^-40^ to minimise erroneous matches due to chance [21,24], (ii) using custom reference databases for mapping and processing reads [23], or (iii) using existing programs designed for short reads, like VSEARCH [39] or CD-HIT [40] with more relaxed settings for clustering error-prone nanopore reads [26,41].

We expect that nanopore metabarcoding studies will become more common, given the release of new nanopore metabarcoding workflows like ASHURE [19], decona [20] and MSI [25], as well improvement of flow cell chemistries and base calling models over time. The latter is evidenced in the decreasing raw read error rate to ∼6% using R9.4 flow cell chemistry [42], and even lower at ∼4% for R10.3 flow cells [43]. In fact, two research groups have independently confirmed that it is now possible to generate highly-accurate, Illumina-like, DNA barcodes without the need for further error correction with R10.3 sequencing chemistry [44,45].

In light of these vast improvements in sequencing accuracy, we propose that the time is ripe for broader-scale nanopore metabarcoding, and on more complex biological communities. In this study, we show that nanopore metabarcoding can produce highly-accurate consensus metabarcodes that are almost always indel-free using R10.3 nanopore reads when processed with the recently-released amplicon_sorter [44]. We further demonstrate that such high-quality metabarcodes can be obtained without the need for complicated wet-laboratory procedures like rolling circle amplification as with the ASHURE workflow, or even error correction, like in the MSI and decona pipelines. Our approach was to metabarcode species-rich, bulk zooplankton samples collected from the tropical waters of Singapore, and benchmark the community composition and relative abundance of nanopore results against conventional Illumina sequencing (Fig. 1). Our study demonstrates the viability of nanopore metabarcoding on complex, biodiverse communities, and we hope this inspires greater confidence in nanopore sequencing for a greater variety of metabarcoding applications.

**Figure 1.**
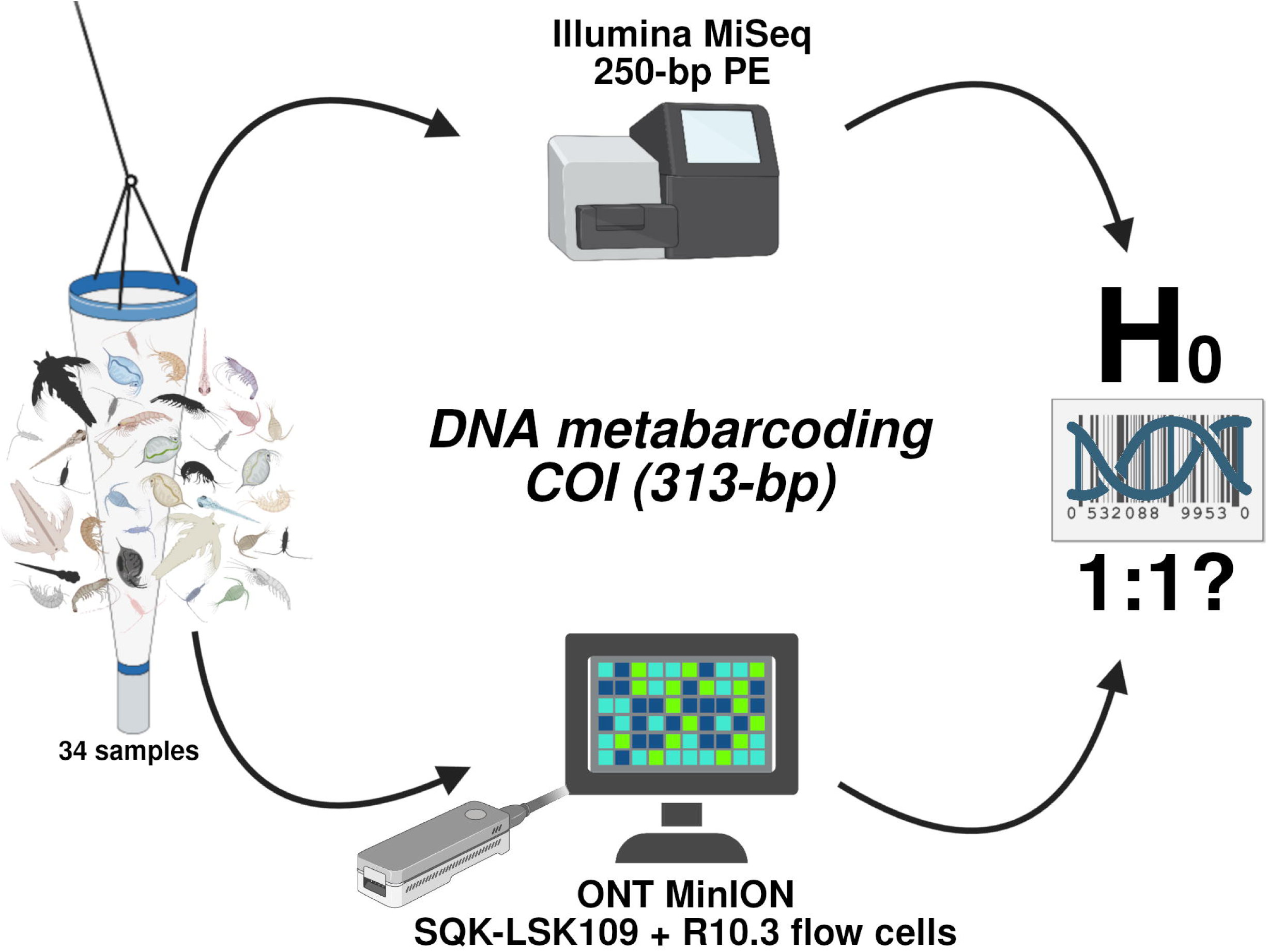
Study workflow from sample collection to sequencing on Illumina and nanopore, with the main aim of determining if nanopore-based DNA metabarcoding would produce comparable results with that of conventional Illumina DNA metabarcoding. Icons adapted from BioRender.

## Methods and Analyses

### Sample collection and processing

The study samples comprised a series of zooplankton collections made during August–September 2020 in Singapore. Collections were permitted by the National Parks Board, Singapore (Permit Number NP/RP18-051). The targeted sites were off Pulau Hantu and Sisters’ Islands in the Singapore Strait (See Supplementary File S1 for GPS coordinates). All plankton collections were performed at night (1800–2200 hrs), and sampling was conducted in two ways. First, triplicate oblique plankton tows were performed from a boat with bongo nets (2 m in length, 500 µm mesh size, 50 cm ring diameter) from a depth of 15 m to the surface at 1 m/s. The plankton net was always rinsed with fresh water before each tow, and its contents were collected as the field negative control. After each tow, the contents from one cod-end were poured through 2 mm and 500 µm sieves to filter excess seawater before bulk preservation in molecular-grade ethanol. Specimens larger than 1 cm were picked out individually. The collected material were thus separated into three size fractions—1 cm, 2 mm and 500 µm. Second, a quatrefoil light trap (30 cm diameter by 25 cm height; 5 mm entry slit width) fitted with two GT-AAAs (Glo-Toob) was left at the jetty of each island 1.5 m below the water surface for two hours (See Supplementary File S1 for GPS coordinates). Light trap samples were processed in the same way as bongo net samples. All bulk samples were brought back to the laboratory and stored at -20 °C prior to DNA extraction.

### DNA extraction and PCR amplification

Bulk samples were first ground with pre-sterilized mortar and pestles. Genomic extraction was performed with DNeasy Blood and Tissue Kit (Qiagen) following the manufacturer’s protocol, except that genomic DNA was eluted in nuclease-free water. To prevent cross-contamination, a fresh set of autoclaved mortar and pestle was used for each tow/light trap. All units were thoroughly washed and autoclaved before the next set of DNA extractions.

We amplified the 313-bp fragment of the mitochondrial cytochrome oxidase c subunit I (COI) gene so that the PCR products could be directly compared across both sequencing platforms (Fig. 1). PCR amplification was performed using the mlCOIintF: 5’-GGW ACW GGW TGA ACW GTW TAY CCY CC-3’ [46] and LoboR1: 5’-TAA ACY TCW GGR TGW CCR AAR AAY CA-3’ [47] primer combination. This primer combination was also chosen for its high amplification success in marine organisms [48–51], and is approximately four times cheaper than the conventional mlCOIintF and jgHCO2198 [52] metabarcoding primer pair [26,53]. Furthermore, Yeo et al. [54] have also demonstrated that 313-bp COI sequences performed just as well as 658-bp barcodes for species-level identification. The primers were tagged at the 5’ end with custom 13-bp sequences (i.e., “tags”) from Srivathsan et al. [38] to allow for downstream demultiplexing of sequence reads to samples. The longer-than-usual tag lengths were necessary to accommodate the error profile of Kit 9 and R10.3 sequencing chemistry [38] (though it was recently reported that shorter 9-bp tags work well for R10.4.1 sequencing kits and flow cells [55]). Each PCR was assigned its own unique forward and reverse tag combination where possible, and if there were overlapping tag combinations, we separated them into different library pools (i.e., Plate A and B).

PCR was carried out in 25 µl triplicate reactions using 2 µl genomic DNA (100× dilution of original extract), 12.5 µl of GoTaq Green Master Mix (Promega), 2 µl of 10 µM 13-bp tagged forward and reverse primers, 1 µl of bovine serum albumin (1 mg/ml; New England Biolabs) and 7.5 µl of nuclease-free water. A step-up thermocycling profile was used: 1 min denaturation at 94 °C; 5 cycles of 30 sec at 94 °C; 2 min at 45 °C; 1 min at 72 °C; 30 cycles of 30 sec at 94 °C; 2 min at 55 °C; 1 min at 72 °C and a final extension of 3 min at 72 °C. All PCR products were screened on 2% agarose gels stained with GelRed (Biotium Inc.) to ensure appropriate amplification. PCR amplicons were subsequently combined by plate into two pools and purified with SureClean Plus (Bioline). Plate A and B had 48 and 72 amplicons (including negatives and controls) respectively. In total, 34 samples, four field controls, and two PCR negatives were carried forward for Illumina and nanopore library preparation (40 IZ 3 PCRs = 120 amplicons).

### Illumina metabarcoding and bioinformatics

We prepared two Illumina libraries using NEBNext Ultra II DNA Library Prep Kit (New England Biolabs) following the manufacturer’s protocol, up till the adapter ligation step (i.e., PCR-free libraries). Libraries were multiplexed using TruSeq CD Dual Indexes (Illumina). Cleanups were performed using 1.0× AMPure XP beads (Beckman Coulter). The two libraries were pooled together and outsourced for sequencing on a single Illumina MiSeq (2IZ250-bp) lane at the Genome Institute of Singapore.

Illumina reads were processed according to a modified metabarcoding pipeline from Sze et al. [56] and Ip et al. [57]. First, Illumina paired-end reads were merged using PEAR v0.9.6 [58]. Thereafter, OBITools v1.2.13 [33] was used for downstream processing of assembled reads. Specifically, the ngsfilter module was used to demultiplex reads to respective PCR replicates under default settings, where up to 2-bp mismatch was allowed for primer sequences, but no mismatch allowed for tag sequences. Sequence reads were then dereplicated and sorted to samples with obiuniq and obisubset respectively. We retained sequences with ≥5 counts and between 303- and 323-bp in length using obigrep. Subsequently, the filtered reads were further collapsed with obiclean, where sequences with 1-bp difference from each other were considered sequencing errors and further collapsed, and only reads with ‘head’ status were retained. We then concatenated all sequences across all samples, and ran cd-hit-est v.4.8.1 [40] to collapse 100% identical sequences. Any sequence that clustered with PCR negatives or control samples at 100% were eliminated.

### Nanopore metabarcoding and bioinformatics

The same cleaned amplicon pools were used to prepare two nanopore libraries with the Ligation Sequencing Kit (SQK-LSK109) following the manufacturer’s protocol, but end-repair and adapter ligation times were increased to 60 and 15 min respectively [53]. Cleanups were likewise done using 0.9× AMPure XP beads (Beckman Coulter) and the supplied Short Fragment Buffer (SFB). Finally, the two libraries were each sequenced on fresh R10.3 MinION flow cells on MinKNOW v20.10.3 for Ubuntu 16. The R10.3 flow cell chemistry was selected given its improved accuracy and homopolymer resolution [45,59]. The first run (RUN A) lasted 20 h and 30 min, while the second (RUN B) lasted 41 h.

Raw fast5 reads were exported to the National University of Singapore’s High Performance Computing (NUS HPC) Volta cluster for GPU basecalling on NVIDIA Tesla V100 SXM2 32GB with Guppy v5.0.14+8f53ee9, using the super accurate (SUP) model at default settings. We then performed a length filter with NanoFilt v2.8.0 [60] to retain only sequence reads ≥250-bp in length. Subsequently, the sequences were distributed to respective PCR replicates with the demultiplexing module of ONTbarcoder v0.1.9 [45]. We set 313-bp as the read length threshold, and kept the other settings as default. Only sequences deviating up to 2 bp from the tag sequence were accepted in the demultiplexing process, which was possible as tags were designed to differ by ≥3-bp from each other [38]. Moreover, ONTbarcoder recognises and splits self-ligated reads during demultiplexing, thereby retaining more reads for downstream analysis. Thereafter, we concatenated the reads by sample.

For metabarcoding analysis, we used the recently released amplicon_sorter v2022-03-28 [44] to sort and group the nanopore reads based on length and sequence similarity in order to generate consensus metabarcodes. We selected it for two reasons. Firstly, amplicon_sorter performs reference-free clustering which is extremely useful in our case since we did not have a priori knowledge of the community composition of our zooplankton samples. Secondly, amplicon_sorter considers all possible clusters when generating consensus sequences, meaning it can be utilised to analyse DNA metabarcoding data. We adopted a conservative approach where sequences were added into a species group by the program only if they were ≥97% similar (--similar_species), and consensus sequences were combined together only if they were ≥98% similar (--similar_consensus). We also set the minimum and maximum length limits to 293- and 333-bp respectively, and performed 3× random sampling (--maxreads) to increase likelihood of sampling rare reads. We then mapped the sequences of each cluster back to the respective consensus sequence with minimap2 v2.24 [61] and polished the consensus with medaka v1.7.2 (https://github.com/nanoporetech/medaka), using the r103_sup_g507 model. Finally, we removed sequences that were present in our PCR negatives and controls from the samples using the same method described for Illumina metabarcoding.

### MOTU delimitation and community analysis

We concatenated both Illumina and nanopore datasets together and aligned the sequences with MAFFT v.7.487 [62], before grouping them into molecular operational taxonomic units (MOTUs) with objective clustering (https://github.com/asrivathsan/obj_cluster) at the 3% threshold. This was consistent with distance thresholds applied in past studies [48,57,63]. We then ran blastn (e-value: 1e^-6^ and 80% identity) with BLAST+ v2.12.0 [64] against the NCBI *nt* database (downloaded 13 June 2022), and obtained taxonomic identities for blast hits that had 85% identity match and minimum 250-bp overlap with readsidentifier v1.1.2 [65]. With the taxa identified, we grouped our Illumina MOTUs for a translation check on Geneious Prime v2022.2.2. (http://www.geneious.com/), using codes 2 (Chordata), 4 (Cnidaria), 5 (all other invertebrates), 9 (Echinodermata and Rhabditophora) and 13 (Ascidiacea). Illumina sequences that failed the translation check were considered possible nuclear mitochondrial DNA (NUMT) and discarded. For MOTUs that matched at ≥90%, we also screened the taxonomic identities against World Register of Marine Species (WoRMS; downloaded 8 May 2022) to confirm the MOTUs were marine, and also against past studies [48,57,59,63,66,67] as well as SeaLifeBase (https://www.sealifebase.ca/) to confirm the MOTUs’ geographic ranges were within the Indo-Pacific.

With the final consolidated MOTU dataset, we assessed if and how MOTU communities compared between sequencing types quantitatively using diversity metrics, permutational multivariate analysis of variance (PERMANOVA), and qualitatively by examining the agreement in MOTU composition in terms of proportion and abundance. All statistical analyses were performed in R v4.3.1 [68], in RStudio (build 2023.03.0) unless otherwise stated, and all relevant plots were generated with the ggplot2 v3.4.2 package [69]. We computed the MOTU richness, Shannon-Wiener, and Simpson indices for each sequencing dataset using the diversity function in vegan v2.6-4 [70] and ran a paired, nonparametric Wilcoxon signed-rank test to test whether differences in the indices were due to different sequencing platforms. We also plotted the rarefaction curves of MOTU richness for each dataset with iNEXT v3.0.0 [71] to examine the relationship between MOTU richness and sampling depth. Community similarities between sequencing types were assessed using: (i) the Jaccard similarity coefficient by converting the MOTU community matrix to binary absence/presence data; and (ii) also with Bray-Curtis distances, where we normalised our MOTUs by relative abundance of sequencing reads [72]. We visualised the distances using nonmetric multidimensional scaling (nMDS) plots (metaMDS in vegan) and heatmaps constructed with pheatmap v1.0.12 package [73]. We also performed PERMANOVA with adonis2 in vegan to test for community differences between Illumina and nanopore sequencing. Here, sequencing type (Illumina or nanopore) was included as a variable, in addition to site (Pulau Hantu or Sisters’ Islands), date (5 August 2020, 19 August 2020, 20 August 2020, 2 September 2020, 3 September 2020 or 16 September 2020), as well as fraction (1 cm, 2 mm or 500 µm). We first verified that each variable had a non-significant betadisper result before inclusion into PERMANOVA. We also analysed the datasets separately to confirm the same ecological conclusions would be obtained regardless of sequencing type. For this, we used the same Bray-Curtis distance datasets, and visualised the community dissimilarities with nMDS. For PERMANOVA, we only incorporated the bongo net samples as that sampling method had the most samples. We used the same three variables (site, date, fraction) and groupings as above for PERMANOVA with adonis2.

We also examined MOTU community compositions to determine how consistent they were between nanopore and Illumina platforms. We first looked at MOTU composition based on phyla, and compared the relative proportions of each phylum at the sequencing dataset level, and further at the sample level. In addition, we were also interested to know if a MOTU that was abundant in nanopore sequencing would be similarly so with Illumina sequencing. For each sample, we sorted and ranked the MOTUs by sequencing reads, and then assessed similarity in rank order of MOTUs between sequencing platforms with Kendall rank correlation coefficient (Kendall’s τ) [74]. We performed the correlation analysis only for 31 out of 34 samples as the remaining three samples had only one pairwise comparison.

### Sequencing accuracy and quality of nanopore reads

A known drawback of nanopore sequencing is its relatively high error rates. A close examination of the error rates of the raw reads and consensus sequences here was thus necessary to address existing concerns regarding its use. We mapped the nanopore sequences against the cleaned Illumina sequences at the sample-level (e.g., ZPT005 nanopore reads to ZPT005 Illumina reads) with mapPacBio.sh v38.96 in BBTools (script was also recommended for nanopore data; https://sourceforge.net/projects/bbmap/). We maximised mapping sensitivity with the --vslow flag, and mapped two datasets: (i) the demultiplexed reads from ONTbarcoder to estimate raw read error rates and (ii) consensus sequences generated from amplicon_sorter to assess consensus sequence quality. We only considered mappings where the nanopore queries had ≥90% identity match to the Illumina reference sequences, and computed the total error rates, which took into account substitutions, insertions, deletions and ambiguous bases.

Additionally, for each MOTU shared between Illumina and nanopore datasets, we further compared the constituent Illumina and nanopore member sequences of that MOTU with dnadiff v1.3 [75]. As our Illumina sequences were already confirmed to be translatable, and are thus free of frameshift errors and unlikely NUMTs, this comparison allowed us to assess the frequency of indel errors in our nanopore consensus sequences.

### Time sampling of nanopore reads

Given the real-time sequencing properties of the MinION, we also preliminarily examined the relationship between sequencing run time and its effect on the nanopore metabarcoding. It was previously observed that 80–90% of DNA barcodes were obtained within the first few hours of sequencing [37,45] for DNA barcoding studies. Here, we tested if the observed trends would be similar in a nanopore metabarcoding context. We subsampled the nanopore reads generated from each run for every hour for the first three hours of sequencing, followed by every three hours thereafter, until 18 h for RUN A and 39 h for RUN B. For each time period, we repeated the entire workflow from Guppy basecalling to amplicon_sorter (see section ‘*Nanopore metabarcoding and bioinformatics*’). For each time point, we noted down (i) the number of raw reads generated, (ii) the number of reads demultiplexed by ONTbarcoder, and (iii) the number of metazoan MOTUs obtained for each time series dataset.

## Results

### Zooplankton collections

A total of 49 bulk zooplankton samples—24 and 25 from Pulau Hantu and Sisters’ Islands respectively—were collected and included in this study (Supplementary File S1). Of the 49 samples, 37 were bongo net samples, seven were light trap samples, and five were field control samples. After sieving and sorting, the 500 µm size fraction was the most common (29 samples), followed by 2 mm (18 samples), with the 1 cm fraction class having the least (2 samples). PCR amplification was successful for 34 samples (28 bongo net and 6 light trap samples), and nanopore and Illumina libraries were prepared for a total of 40 samples for this comparative study (including four field controls and two PCR negatives).

### Metabarcoding and MOTU delimitation

The two MinION sequencing runs generated 20,045,167 raw reads in total, while Illumina MiSeq sequencing generated 10,038,735 paired-end reads. For Illumina, 7,630,728 reads were successfully assembled with PEAR, 4,218,977 reads were successfully demultiplexed, and 4,162,498 reads remained after the length filter. We obtained 10,788 clean haplotypes after removing sequences present in controls and PCR negatives. For nanopore sequencing, we retained 14,123,752 reads after Guppy basecalling and NanoFilt, and 6,918,618 reads after demultiplexing with ONTbarcoder (48.6% demultiplexing success). Consensus calling with amplicon_sorter generated a total of 4,206 sequences from 3,525,077 reads (51% of demultiplexed reads). At the sample level, 57.6% of demultiplexed reads were utilised by the program to generate consensus sequences on average, with a minimum of 47.1% to 73.3% maximum. The median length was 313-bp (62% of total sequences generated); minimum and maximum sequence lengths were 300- and 339-bp respectively. We also observed that amplicon_sorter very rarely generated consensus sequences from different “gene groups” (two samples had one consensus sequence each while only one sample had five such consensus sequences). These were found to be of non-mitochondrial origin when we conducted nucleotide BLAST searches on NCBI web servers, and were thus excluded from the dataset. After filtering sequences present in the negatives and controls, we retained 3,973 consensus sequences (3,295,247 reads). As polishing with medaka had a minimal impact in reducing error rates (∼0.02% decrease), we carried out the analysis using the unpolished dataset instead (see Vierstraete and Braeckman, [44]).

From the combined sequencing dataset, we obtained 1,031 MOTUs at the 3% threshold, with only 688 identified (at 85% identity match with ≥250-bp overlap) via readsidentifier. We discarded 61 MOTUs as four of them matched only to unclassified environmental samples, while the remainder were Rhodophyta (35 MOTUs), followed by Fungi (10 MOTUs) and Bacillarophyta (8 MOTUs). We further eliminated one Illumina MOTU for failing the translation check, and 10 MOTUs that matched non-marine Insecta. None of the remaining MOTUs’ geographic ranges fell outside the Indo-Pacific. Our final dataset comprised 616 Metazoa MOTUs, of which 316 had ≥97% match to a sequence on NCBI *nt* database, and 274 out of 316 obtained a species-level identity (Supplementary File S1).

### Comparing nanopore and Illumina metabarcoding

The general trend across both sequencing platforms was that if a sample had a higher number of demultiplexed reads compared to other samples within Illumina, the same was true for nanopore, although the differences were greater for the latter (Fig. 2a). Illumina also recovered a higher number of MOTUs (589 vs. 471) than nanopore, but species accumulation curves suggested that ∼120 samples were needed to fully capture zooplankton diversity for both sequencing types (Fig. 2b). 444 MOTUs were shared (72% overlap) across both sequencing platforms, with more MOTUs unique to Illumina than to nanopore (Fig. 2b, insert). At the sample-level, Illumina metabarcoding also consistently recovered more MOTUs than nanopore, with the exception of ZPT017 and ZPT023 (Fig. 2c). MOTU richness (*p*-value = 4.056 x 10^-5^) and Shannon-Wiener diversity (*p*-value = 0.03) were found to be significantly different across paired samples, while Simpson diversity was not (*p*-value = 0.63, Fig. 2d). Even so, we observed clustering by sample on the nMDS plots, especially when using the Bray-Curtis distance metric (Fig. 3). This suggested that although MOTU richness differed across paired samples, the relative abundance of MOTUs within each sample were quite similar across both sequencing platforms. PERMANOVA revealed significant differences in communities for both Jaccard and Bray-Curtis datasets (Jaccard: *df* = 27, *F* = 1.2329, *R^2^* = 0.4542, *p* = 0.0014; Bray-Curtis: *df* = 27, *F* = 1.6542, *R^2^* = 0.52754, *p* = 0.0001), but the differences were driven by the other three variables and not sequencing type (Table 1). When each sequencing dataset was analysed separately, we noted the same ecological conclusions from the nMDS plots and PERMANOVA as well—that the bongo net zooplankton communities were structured by date, fraction and site regardless of the sequencing platform (Fig. 4 and Table 2).

**Figure 2.**
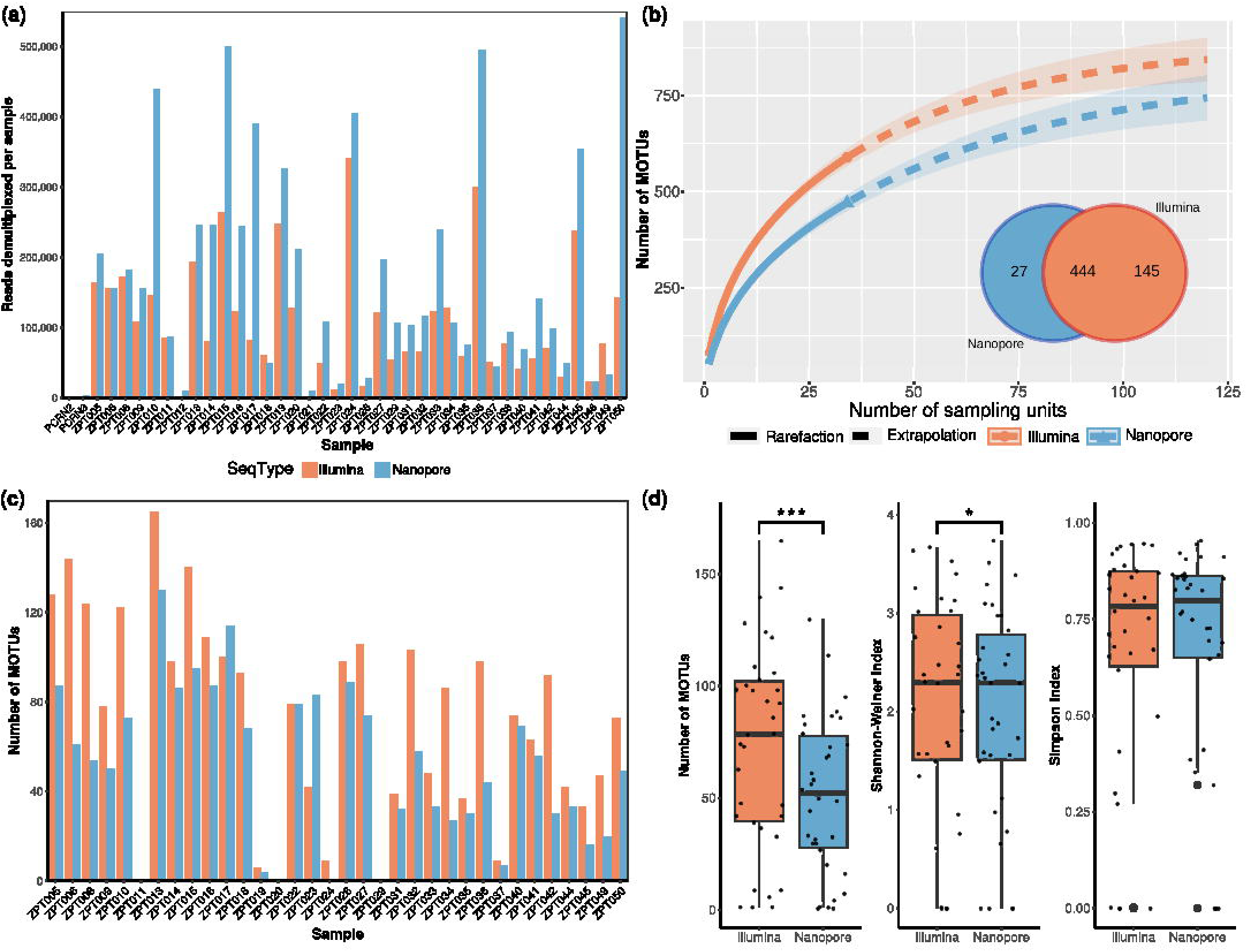
Sequencing statistics of zooplankton metabarcoding with Illumina MiSeq and Nanopore MinION. (a) Bar plot of sequence reads demultiplexed per sample per sequencing dataset. (b) Species accumulation curves of molecular operational taxonomic unit (MOTU) richness for each sequencing platform against the number of samples, extrapolated to visualise number of samples needed to capture maximum richness; number of MOTUs obtained (and shared) expressed in Venn (insert). (c) Bar plots showing the number of MOTUs obtained per sample per sequencing type. (d) Box plots comparing MOTU richness, Simpson index, and Shannon-Weiner index between sequencing platforms; asterisks indicate significant differences for paired Wilcoxon signed-rank tests, and dots represent individual sample points (jittered).

**Figure 3.**
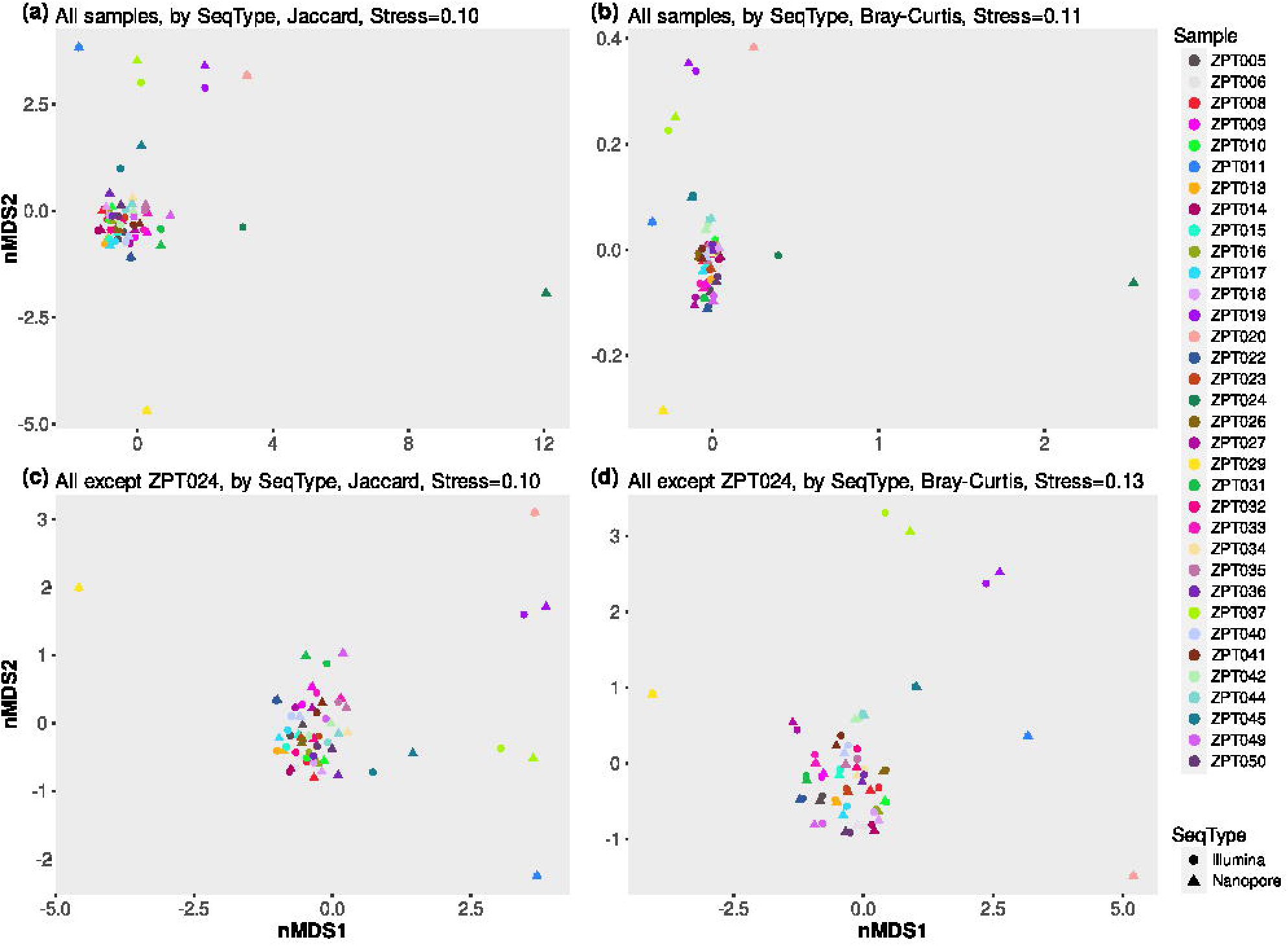
Two-dimensional nonmetric multidimensional scaling (nMDS) plots of Illumina (circles) and nanopore (triangles) community datasets based on Jaccard (a and c) and Bray-Curtis (b and d) distances, coloured by sample, with all samples (a and b), and ZPT024 removed (c and d) to better visualise the spread of points.

**Figure 4.**
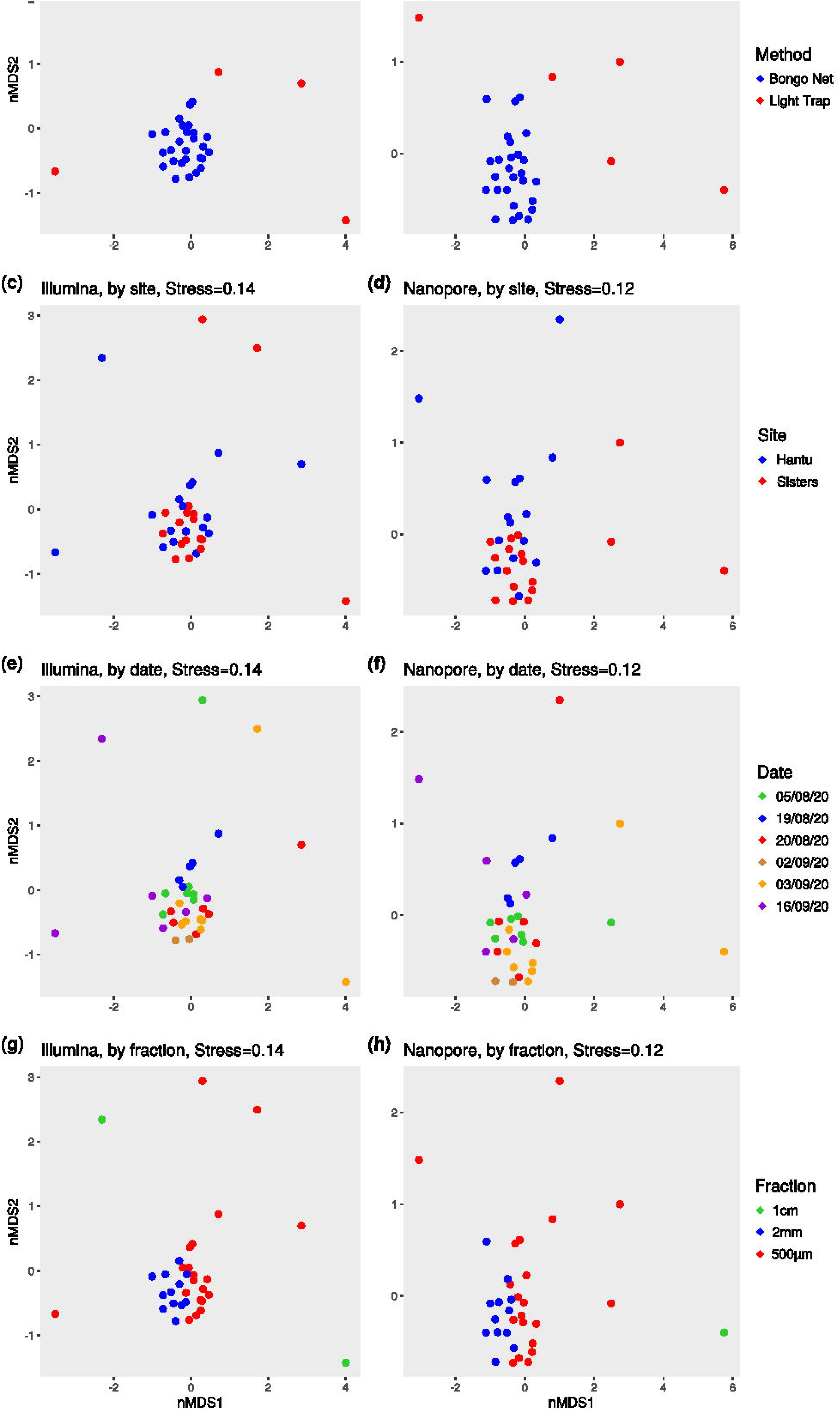
Two-dimensional nonmetric multidimensional scaling (nMDS) plots based on normalised Bray-Curtis distances for Illumina (a, c, e, g), and nanopore (b, d, f, h); coloured by sampling method (a and b), sampling site (c and d), date (e and f), and size fraction (g and h). ZPT024 was removed from the nanopore dataset to better visualise the spread of points; it was similarly distinct from the remaining samples for both sequencing types.

**Table 1.** Permutational multivariate analysis of variance (PERMANOVA) results comparing community differences between nanopore and Illumina metabarcoding datasets, with Jaccard coefficient and Bray-Curtis dissimilarity. Variables with significant *p*-values are highlighted in bold.

Jaccard
| | $df$ | Sum of squares | $R^2$ | F-value | $p$-value |
| --- | --- | --- | --- | --- | --- |
| SeqType | 1 | 0.2065 | 0.00827 | 0.6062 | 0.987 |
| **Site** | **1** | **0.9791** | **0.03922** | **2.8744** | **0.001** |
| **Date** | **4** | **3.0335** | **0.12151** | **2.2264** | **0.001** |
| **Fraction** | **2** | **2.4961** | **0.09999** | **3.6639** | **0.001** |
| SeqType:Site | 1 | 0.0783 | 0.00314 | 0.2298 | 1.000 |
| SeqType:Date | 4 | 0.4419 | 0.01770 | 0.3244 | 1.000 |
| SeqType:Fraction | 2 | 0.2286 | 0.00916 | 0.3355 | 1.000 |
| **Site:Fraction** | **2** | **1.3542** | **0.05424** | **1.9877** | **0.001** |
| **Date:Fraction** | **4** | **2.0055** | **0.08034** | **1.4719** | **0.001** |
| SeqType:Site:Fraction | 2 | 0.2235 | 0.00895 | 0.3281 | 1.000 |
| SeqType:Date:Fraction | 4 | 0.2916 | 0.01168 | 0.2140 | 1.000 |
| Residual | 40 | 13.6253 | 0.54580 |  |  |
| Total | 67 | 24.9641 | 1.00000 |  |  |

Bray-Curtis
| | $df$ | Sum of squares | $R^2$ | F-value | $p$-value |
| --- | --- | --- | --- | --- | --- |
| SeqType | 1 | 0.0607 | 0.00263 | 0.2231 | 0.999 |
| **Site** | **1** | **1.2093** | **0.05248** | **4.4435** | **0.001** |
| **Date** | **4** | **4.0874** | **0.1774** | **3.7547** | **0.001** |
| **Fraction** | **2** | **3.7336** | **0.16204** | **6.8594** | **0.001** |
| SeqType:Site | 1 | -0.0012 | -0.00005 | -0.0046 | 1.000 |
| SeqType:Date | 4 | 0.0419 | 0.00182 | 0.0385 | 1.000 |
| SeqType:Fraction | 2 | 0.0197 | 0.00085 | 0.0361 | 1.000 |
| **Site:Fraction** | **2** | **1.6076** | **0.06977** | **2.9535** | **0.001** |
| Date:Fraction | 4 | 1.3671 | 0.05933 | 1.2559 | 0.104 |
| SeqType:Site:Fraction | 2 | 0.0016 | 0.00007 | 0.0030 | 1.000 |
| SeqType:Date:Fraction | 4 | 0.0274 | 0.00119 | 0.0252 | 1.000 |
| Residual | 40 | 10.886 | 0.47246 |  |  |
| Total | 67 | 23.0409 | 1.00000 |  |  |

**Table 2.** Permutational multivariate analysis of variance (PERMANOVA) results comparing bongo net communities for nanopore and Illumina datasets, using Bray-Curtis dissimilarity. Variables with significant *p*-values are highlighted in bold.

Nanopore
| | $df$ | Sum of squares | $R^2$ | F-value | $p$-value |
| --- | --- | --- | --- | --- | --- |
| **Site** | **1** | **0.5841** | **0.08491** | **3.8896** | **0.001** |
| **Date** | **4** | **1.8744** | **0.27248** | **3.1203** | **0.001** |
| **Fraction** | **1** | **1.2382** | **0.18000** | **8.2451** | **0.001** |
| Site:Fraction | 1 | 0.2479 | 0.03604 | 1.6510 | 0.090 |
| Date:Fraction | 4 | 0.6818 | 0.09911 | 1.1350 | 0.286 |
| Residual | 15 | 2.2526 | 0.32746 |  |  |
| Total | 26 | 6.8791 | 1.00000 |  |  |

Illumina
| | $df$ | Sum of squares | $R^2$ | F-value | $p$-value |
| --- | --- | --- | --- | --- | --- |
| **Site** | **1** | **0.5412** | **0.07609** | **3.3736** | **0.001** |
| **Date** | **4** | **2.0928** | **0.29420** | **3.2611** | **0.001** |
| **Fraction** | **1** | **1.1031** | **0.15508** | **6.8759** | **0.001** |
| Site:Fraction | 1 | 0.2723 | 0.03828 | 1.6973 | 0.082 |
| Date:Fraction | 1 | 0.6974 | 0.09803 | 1.0867 | 0.354 |
| Residual | 15 | 2.4065 | 0.33831 |  |  |
| Total | 26 | 7.1133 | 1.00000 |  |  |

**Table 3.** Pairwise comparison of the top 10 (where possible) most abundant molecular operational taxonomic units (MOTUs) within each sample, by read counts, from each sequencing type. The MOTU could either be found in the same rank order across both sequencing types [Scenario 1], or within the top *n* number of comparisons made, but not in the same rank order [Scenario 2], or present in the top *n* in one sequencing dataset, but not the other [Scenario 3]. Percentages are included in round parenthesis.

| Sample | MOTUs compared | Perfect [Scenario 1 only] | Congruent [Scenarios 1+2] |
| --- | --- | --- | --- |
| ZPT005 | 10 | 7 (70%) | 9 (90%) |
| ZPT006 | 10 | 10 (100%) | 10 (100%) |
| ZPT008 | 10 | 2 (20%) | 10 (100%) |
| ZPT009 | 10 | 8 (80%) | 10 (100%) |
| ZPT010 | 10 | 5 (50%) | 9 (90%) |
| ZPT011 | 1 | 1 (100%) | 1 (100%) |
| ZPT013 | 10 | 5 (50%) | 10 (100%) |
| ZPT014 | 10 | 3 (30%) | 9 (90%) |
| ZPT015 | 10 | 8 (80%) | 10 (100%) |
| ZPT016 | 10 | 7 (70%) | 9 (90%) |
| ZPT017 | 10 | 5 (50%) | 8 (80%) |
| ZPT018 | 10 | 3 (30%) | 9 (90%) |
| ZPT019 | 6 | 4 (67%) | 4 (67%) |
| ZPT020 | 1 | 1 (100%) | 1 (100%) |
| ZPT022 | 10 | 2 (20%) | 9 (90%) |
| ZPT023 | 10 | 4 (40%) | 8 (80%) |
| ZPT024 | 9 | 1 (11%) | 1 (11%) |
| ZPT026 | 10 | 3 (30%) | 9 (90%) |
| ZPT027 | 10 | 1 (10%) | 8 (80%) |
| ZPT029 | 1 | 1 (100%) | 1 (100%) |
| ZPT031 | 10 | 7 (70%) | 9 (90%) |
| ZPT032 | 10 | 4 (40%) | 9 (90%) |
| ZPT033 | 10 | 5 (50%) | 10 (100%) |
| ZPT034 | 10 | 8 (80%) | 9 (90%) |
| ZPT035 | 10 | 5 (50%) | 9 (90%) |
| ZPT036 | 10 | 5 (50%) | 9 (90%) |
| ZPT037 | 9 | 7 (78%) | 7 (78%) |
| ZPT040 | 10 | 4 (40%) | 9 (90%) |
| ZPT041 | 10 | 5 (50%) | 9 (90%) |
| ZPT042 | 10 | 4 (40%) | 8 (80%) |
| ZPT044 | 10 | 3 (30%) | 9 (90%) |
| ZPT045 | 10 | 7 (70%) | 9 (90%) |
| ZPT049 | 10 | 5 (50%) | 9 (90%) |
| ZPT050 | 10 | 2 (20%) | 9 (90%) |

Since MOTU richness differed between each sample’s Illumina and nanopore datasets, we checked if this difference altered the respective community compositions. Both Illumina and nanopore recovered all 10 metazoan phyla, with nanopore recovering an additional singleton Platyhelminthes MOTU. Proportions of phyla were found to be consistent across both sequencing datasets, and were largely dominated by Arthropoda (∼53%), followed by Chordata (∼20%) and then Cnidaria (∼12%) (Fig. 5a and Table S1). The differences in MOTU richness were largely from these three dominant groups, with Illumina recovering 1.2 to 1.3× more MOTUs from each of these three phyla compared to nanopore (Table S1). The largest disparity was in Mollusca, for which Illumina recovered twice the number of MOTUs than nanopore. For the remaining six phyla (Echinodermata, Annelida, Porifera, Chaetognatha, Ctenophora, Bryozoa), Illumina and nanopore recovered approximately the same number of MOTUs. At the sample-level, the similar phylum proportions were also consistently observed, albeit with differences in species numbers (Fig. 5b). Only ZPT024 was markedly different in terms of community composition, and this was consistent with the stark dissimilarity observed with nMDS plots (Figs. 3a and b). When ranking the MOTUs by sequencing read counts between sequencing platforms, we found that Kendall’s τ was significantly positive for 30 samples (min: 0.484; max: 0.986; *p*-value << 0.05; Table S2), which suggested a positive correlation in MOTU rank abundance between both sequencing platforms. Kendall’s τ was also positive for ZPT024 (0.478), but the *p*-value was insignificant. This meant that if a MOTU was found to be abundant in one sample for one sequencing dataset, it would be highly likely to be abundant in the alternative platform as well. This assessment corroborated with the high pairwise Bray-Curtis similarity observed between samples across both sequencing platforms (Fig. S1), since the metric took into account read count data. This further demonstrated that nanopore metabarcoding could reliably and consistently recover abundant MOTUs; this was similarly corroborated by [26], even though our bioinformatic pipelines differed.

**Figure 5.**
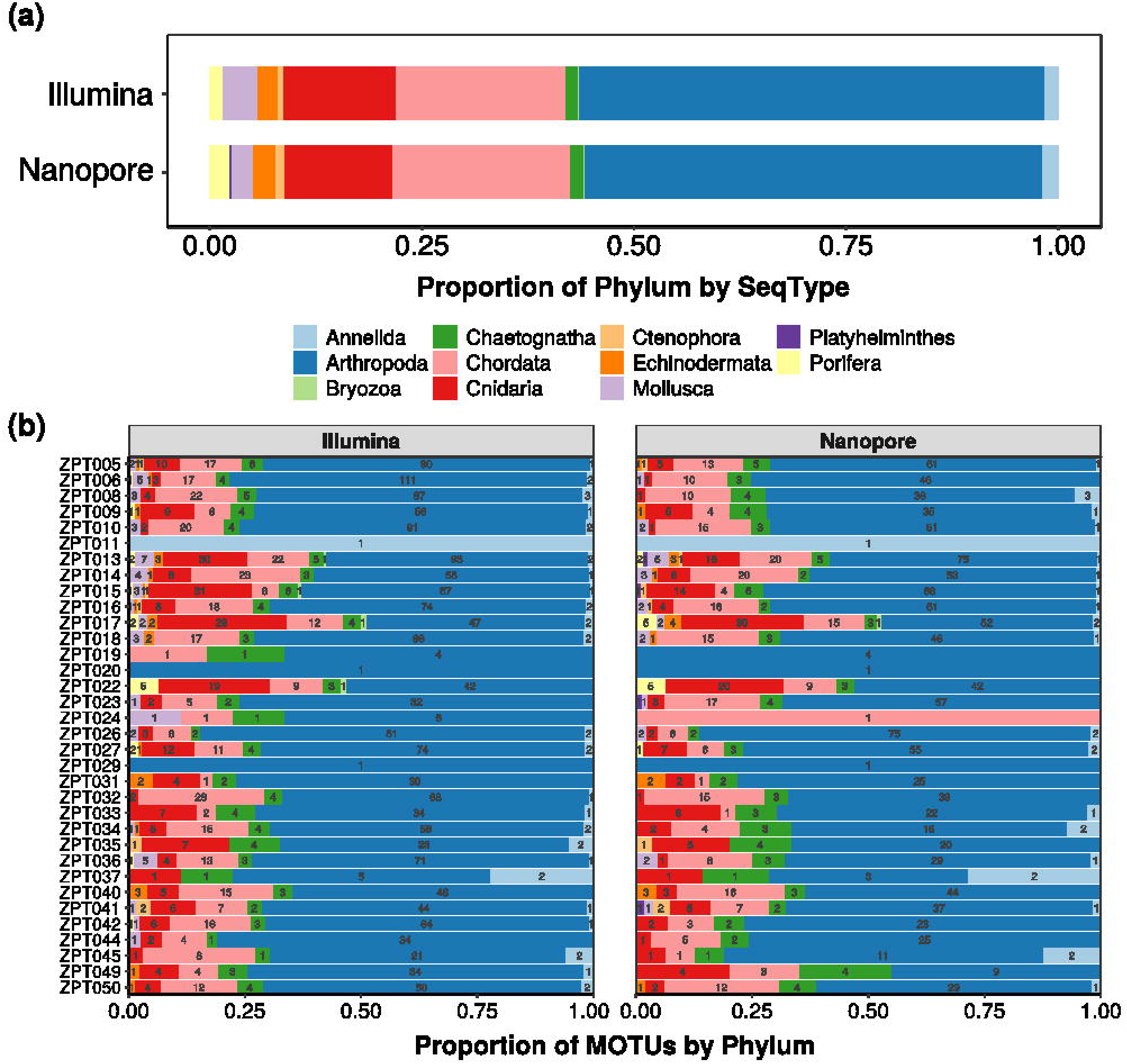
Bar plots showing the relative proportions of molecular operational taxonomic units (MOTUs, grouped by phylum) by sequencing type (a), and by sample (b).

### Nanopore metabarcode quality

We found that ∼98% of the raw nanopore reads were erroneous when mapped to their respective Illumina samples, with a mean error rate of 4.20% (Fig. 6 and Table S3). This was consistent with the 4% error rate reported by Gunter et al. [43] for R10.3 flow cell chemistry. After consensus calling with amplicon_sorter however, and without further polishing with medaka, the percentage of consensus sequences per sample that remained erroneous dropped to 0–50.0% (average 24.0%), and error rates correspondingly decreased to 0–1.18% (average 0.40%) (Fig. 6 and Table S3).

**Figure 6.**
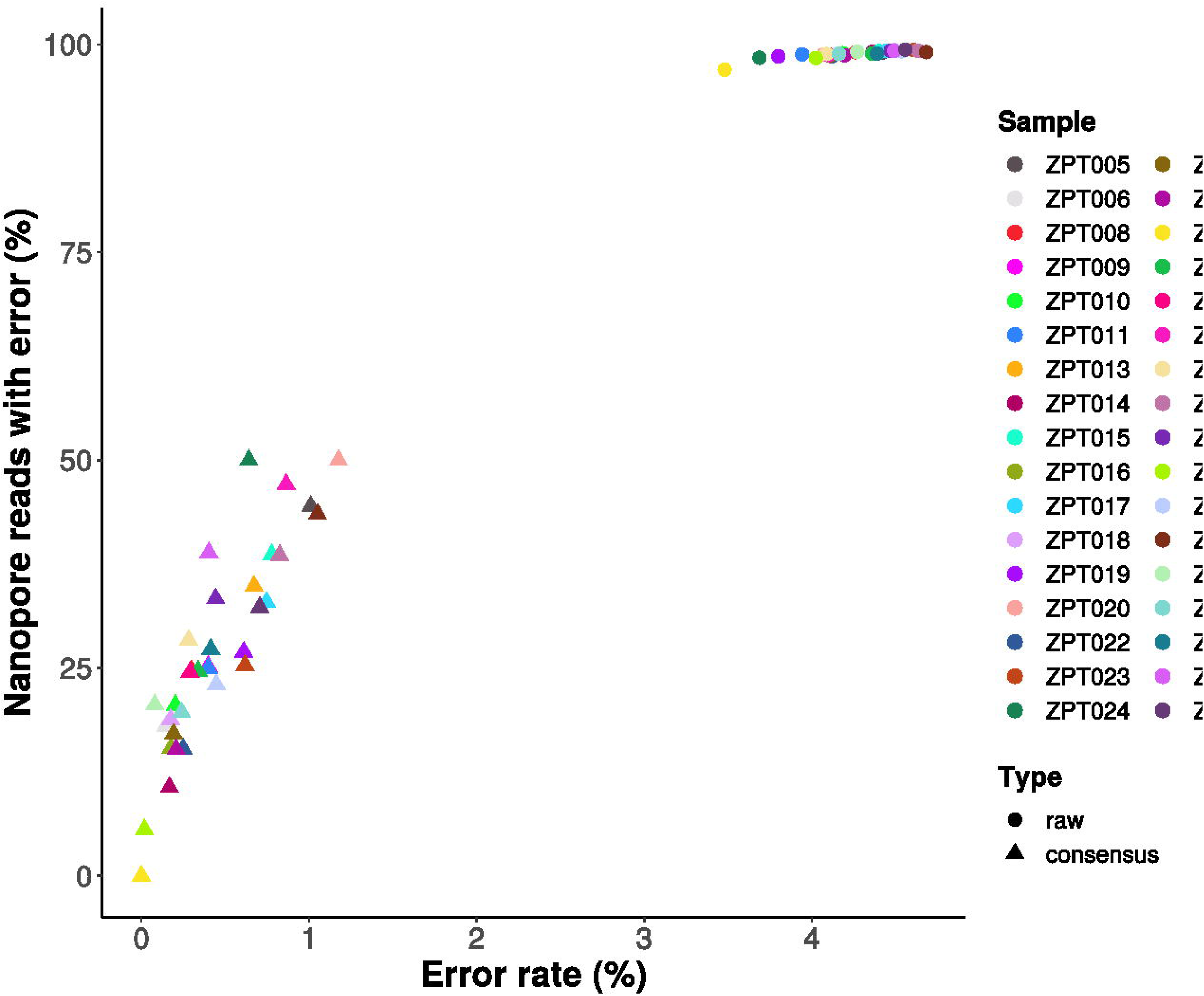
Scatterplot showing the percentage (y-axis) of raw reads (circles) as well as consensus sequences (triangles) of each sample that were erroneous and the corresponding error rates (x-axis), when benchmarked against the respective sample’s Illumina dataset.

Furthermore, for the 444 MOTUs shared between Illumina and nanopore, nanopore sequences from 406 MOTUs (91.4%) did not have indel errors when compared to the same MOTU’s Illumina sequences (Table S4). For the remaining 38 MOTUs: 22 of them had nanopore sequences with 1 indel-error, five with 2 indel errors. The rest had three or more indel errors, but this only affected 11 MOTUs. Since our Illumina sequences were already confirmed to be translatable, it in turn confirmed that 91.4% of the nanopore consensus sequences were free of any frameshift errors, and thus translatable as well.

### Nanopore sequencing with time

We subsampled the fast5 reads of each run for every hour for the first three hours, and every three hours thereafter to investigate the relationship of (i) number of raw reads, (ii) number of demultiplexed reads, and (iii) number of metazoan MOTUs obtained over time. Although the number of samples differed between runs, both runs showed a similar trend in that all three variables increased at a decreasing rate over time (Fig. 7). Raw reads and demultiplexed reads both increased fairly proportionately with respect to each other, with both variables only starting to plateau near the end of the respective runs. Conversely, metazoan MOTUs largely stabilised by the midway mark of each run, with RUN A and B obtaining 85% of the final MOTU count by the 12- and 15-hour mark respectively (Table S5). Beyond that, however, further increase in reads did not translate to substantial increase in metazoan MOTUs.

**Figure 7.**
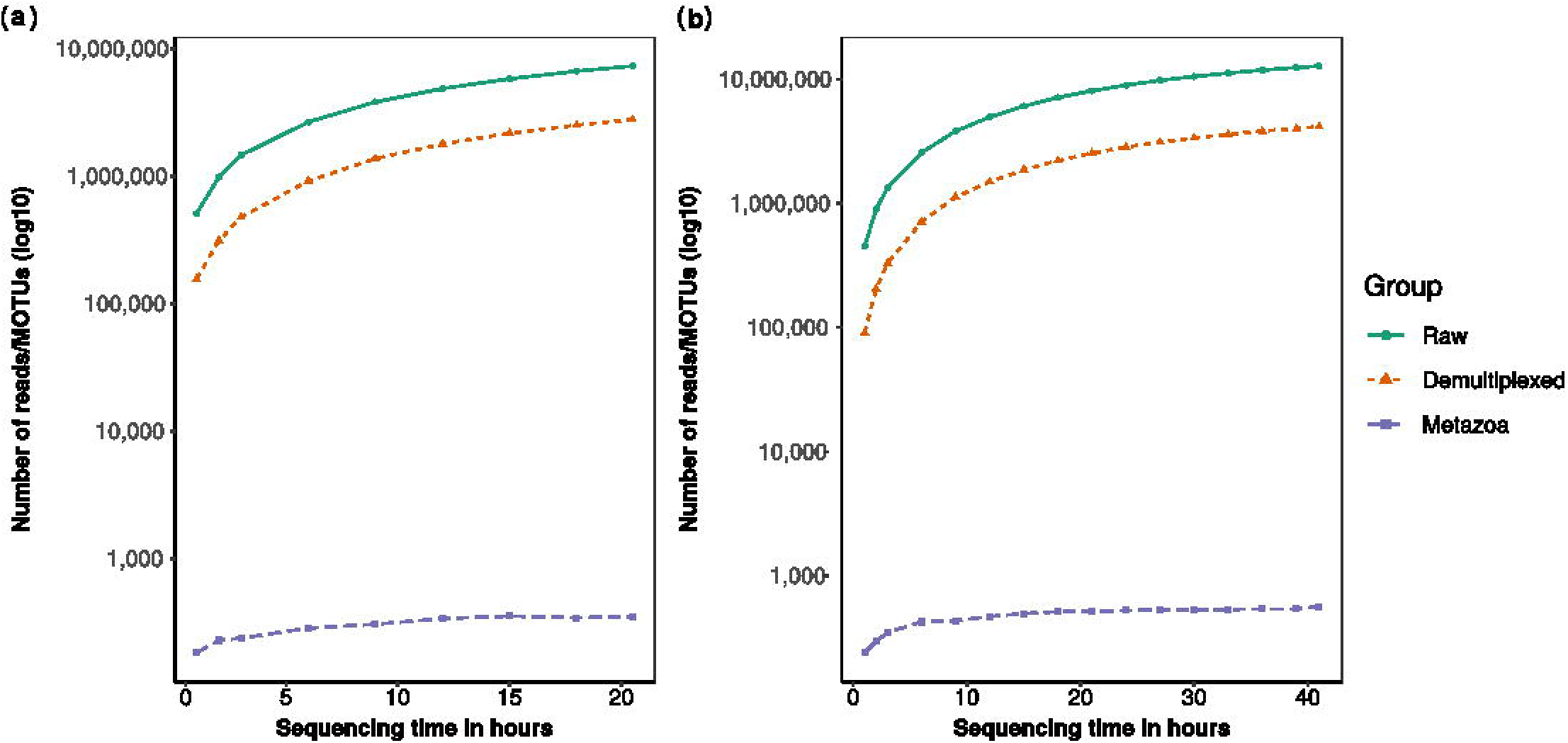
Line graphs showing the change in number of raw reads (green), demultiplexed reads (orange) and metazoan MOTUs (purple) with sequencing run time, for RUN A (left) and RUN B (right).

## Discussion

Using a set of zooplankton samples as our case study, we applied a nanopore-based metabarcoding approach using ONT’s MinION sequencer, and processed the nanopore reads with the recently released amplicon_sorter to show that nanopore metabarcodes are highly accurate and almost always indel-free. We thus demonstrate that nanopore-based metabarcoding is viable, and ready to be incorporated into more projects. We do note that nanopore metabarcoding is not perfect, and we examine the strengths and weaknesses of nanopore metabarcoding with amplicon_sorter.

### Nanopore metabarcodes are highly accurate and virtually indel-free

It is now possible to achieve highly accurate nanopore consensus metabarcodes with amplicon_sorter. In our case, nanopore consensus metabarcodes were observed to be ∼99.6% accurate when benchmarked against their respective Illumina samples. We note this to be slightly better than the median 99.3% sequencing accuracy observed by Baloğlu et al. [19], which could be due to our use of more recent sequencing chemistry and base calling model. Furthermore, amplicon_sorter generated consensus metabarcodes that did not require further polishing, mirroring an initial observation made by Srivathsan et al. [45] that led to algorithmic changes in ONTbarcoder. This is in contrast to prior nanopore metabarcoding pipelines that always included a polishing step, e.g., Egeter et al. [25] polished their sequences with RACON [76], while decona [20] incorporated medaka for polishing. Our results suggest that this is no longer needed, as it only resulted in a negligible 0.02% improvement in error rates for our nanopore metabarcodes. The high level of sequencing accuracy is undoubtedly attributable to the R10.3 flow cell chemistry and its improved ability to resolve homopolymeric regions, in combination with the latest super accurate (SUP) basecalling model released by ONT. This is advantageous as it saves on time and computational resources, because each consensus sequence has to be polished individually when running medaka. For our dataset alone, nearly 4,000 instances of medaka were performed, and this is unlikely to scale well computationally for more diverse, or larger-scale metabarcoding projects, where the number of consensus sequences obtained are expected to increase. An added advantage was that almost all our unpolished nanopore metabarcodes (91.4%) were indel-free when compared to their Illumina counterparts, with nearly all of the 38 remaining nanopore sequences having only 1–2 indel errors. Most of the nanopore sequences can thus be subjected to translation checks without error, which would help to rule out chimaeras while boosting confidence in the quality of obtained nanopore metabarcodes. Furthermore, we were able to achieve all these using the amplicon_sorter program alone with just a single command.

### Lower MOTU richness with nanopore metabarcoding than Illumina

While we have demonstrated that nanopore metabarcoding generated metabarcodes with Illumina-like quality, we recognise that it yielded certain differences in other aspects when benchmarked against Illumina. The most notable difference was in MOTU richness, where we obtained 589 Illumina MOTUs, compared to 471 nanopore MOTUs, with 444 MOTUs shared across both platforms (72% congruence) (Fig. 1b, insert). This was corroborated by a significant difference from the paired Wilcoxon signed-rank test (Fig. 1d). We investigated two potential reasons for this observed difference in MOTU richness between Illumina and nanopore.

The first reason was resolution limits of amplicon_sorter, presently at 95–96% [44]. This means that closely-related species, with less than 4% variance in the COI sequence, will be grouped together by amplicon_sorter, resulting in a lower number of MOTUs obtained. This was challenging to determine as our zooplankton samples were not mock communities, and we did not have prior knowledge of closely-related species groups that we could use to evaluate the resolution limits. We screened ZPT024 and ZPT034—samples that had the lowest Jaccard similarity coefficients between Illumina and nanopore. We first searched for a MOTU that was detected in both Illumina and nanopore for that sample, and then checked if there were any congenerics found in Illumina but not in nanopore (we assumed that congenerics had a higher likelihood of being closely-related compared to other taxonomic ranks). We then checked if the pairwise *p*-distance between these sequences differed by ≤4%, but since we did not encounter any such instance, we do not think that the resolution limit of amplicon_sorter was the main contributing factor for differences in MOTU richness for our study. We emphasise that future users pay special heed to this resolution limit when selecting metabarcoding loci. For instance, zooplankton metabarcoding studies have used hypervariable regions in nuclear 18S rRNA [77–79], nuclear 28S rRNA [80], and mitochondrial 16S rRNA [81] in addition to COI [82–85]. The chosen loci must be divergent enough so that the species groups would not be over-collapsed by amplicon_sorter.

The second potential cause for difference in MOTU richness was based on the observation that since amplicon_sorter grouped only ∼57% of the reads on average for consensus calling, we checked if the MOTUs unique to Illumina could be found in the unsorted nanopore reads. We mapped the ungrouped nanopore reads to the unique Illumina MOTUs with mapPacBio.sh (same settings), and found that had amplicon_sorter incorporated these reads, 22 ZPT samples would have had a complete overlap with the MOTUs detected by Illumina sequencing. The remaining 10 samples would mostly still lack 1–2 MOTU(s), with only ZPT008 and ZPT049 missing four or five MOTUs respectively. We further found that the unsorted nanopore reads had a comparatively higher total error rate of ∼4.52%, above the distance or length thresholds for forming and grouping clusters. This implied that bioinformatic processing of reads by amplicon_sorter was the more likely reason for the MOTU difference. Further tests however, are needed to better optimise consensus calling settings with amplicon_sorter.

In any case, we note that the aforementioned limitations of amplicon_sorter will not pose a major issue to future metabarcoding projects, given that ONT is continuously updating its flow cell chemistry and basecalling algorithms. Its most recent pivot to R10.4.1 flow cell version and v14 kit chemistry (SQK-LSK114) offers Q20+ raw read accuracy (i.e., 1 in 100 error rate). Potential implications would most certainly be higher-quality raw reads that allow for more precise formation and merging of species groups by amplicon_sorter, which in turn will likely improve the resolution limits of the algorithm. For instance, Ni et al. [86] and Sereika et al. [87] have reported ∼99.1% modal raw read accuracy when using the latest R10.4 sequencing chemistry—a considerable improvement compared to the v9 + R10.3 sequencing chemistry we used. It is thus quite foreseeable the limiting factors of amplicon_sorter will resolve as nanopore read quality improves with time.

### Nanopore metabarcoding costs

Various studies have compared sequencing costs between nanopore and Illumina for metabarcoding, and it is generally agreed upon that nanopore metabarcoding with the MinION is generally cheaper than Illumina MiSeq [25,26]. We reduced reagent costs further by adopting a single-PCR tagging strategy, where each of our PCR primers were tagged on 5’-end with 13-bp tags [45]. This enabled us to pool multiple PCR replicates into just two pools for nanopore library preparation without further need to barcode them. The only downside was that it required a separate software (e.g., ONTbarcoder) rather than Guppy for sample demultiplexing. However, the single PCR-tagging saved us processing time because the tagging occurred during thermocycling rather than as an additional step in the library preparation process (thermocycling runs for the same length of time regardless whether tagging is performed). The general utility of tagged-PCR primers also means that it can be used for other DNA sequencing projects [59,63,88], and even for Illumina sequencing (like in this study).

## Potential Implications

### Nanopore metabarcoding for community characterisation

From an operational perspective, we have demonstrated that nanopore-based metabarcoding is viable when benchmarked against Illumina sequencing. Our nanopore metabarcodes were virtually Illumina-like, even with (soon-to-be-obsolete) v9 library preparation kits and R10.3 MinION flow cells. This is only going to improve moving forward, and it is time to relinquish the perception that nanopore sequencing produces highly erroneous reads. Even though there were differences between sequencing platforms, we ultimately found that the same ecological conclusions were obtained regardless—that our zooplankton communities were structured by date, site and fraction, and using a different sequencer was not a significant factor in explaining zooplankton community dissimilarities. Even the relative abundance of MOTUs was fairly consistent across sequencing platforms (88% congruence) and both sequencers successfully recovered 10 metazoan phyla. This also means that future users can employ nanopore sequencing for community metabarcoding with the confidence that their results will be consistent with Illumina, with the potential to leverage the cost-effectiveness, portability and real-time advantages that nanopore sequencing brings. For example, some studies have already incorporated in-situ nanopore metabarcoding on board marine vessels [23,24], and we believe more will follow suit in future, especially in the field of plankton monitoring. We did observe however, that amplicon_sorter was less likely to recover rarer MOTUs in the community compared to Illumina. Hence, users who wish to detect rarer species with degenerate primer sets will have to go with conventional Illumina sequencing in order to increase the chances of detection. We do believe this drawback can be soon addressed given that the latest and most accurate R10.4.1 sequencing chemistry is already available, and there are an increasing number of promising reports regarding its use [55,86,87]. Further benchmarking studies will be needed to investigate how these improvements impact metabarcoding.

### Sequencing turnaround times

Another attractive property of nanopore sequencing is its ability to sequence in real-time. Users can terminate the run when their sequencing needs have been met, wash the flow cell and even recycle it for future use. We were thus interested to know if there was a “sweet-spot” for MOTU richness obtained in relation to sequencing run time for metabarcoding sequencing, based on the observation that up to 90% of DNA barcodes were obtained within the first few hours [45]. Our preliminary examination from subsampling nanopore reads with time was that both runs reached ∼85% of the final MOTU count in under 12 h and 15 h for RUNs A and B respectively (Fig. 7 and Table S5), and sequencing beyond that did not lead to a substantial increase in the number of metazoan MOTUs recovered. We recognise that the relationship between run time and MOTUs recovered is not immediately clear for nanopore metabarcoding (vis-à-vis DNA barcoding). Metabarcoding is likely to be more sensitive to factors such as the number of samples pooled into one flow cell, flow cell health (different flow cells may start with different number of pores available for sequencing) and even pore occupancy (percentage of pores actively sequencing). More tests on the number of metabarcoding samples that can be comfortably multiplexed onto a MinION flow cell without compromising recovered MOTU diversity are needed. What was clear however, was that turnaround times were much faster; it took us three days to complete both nanopore runs (we ran RUN A and B consecutively), in contrast to outsourcing Illumina MiSeq sequencing, which would take ∼2–4 weeks at the very least. Researchers have even taken advantage of this quicker turnaround time in time-sensitive situations such as disease surveillance [89]. Even for zooplankton biomonitoring, where sampling intervals can be as often as every two weeks [90], a nanopore-based metabarcoding approach would enable a quicker generation of results that make proposed routine biomonitoring strategies like Song et al. [91] more operationally feasible.

## Conclusion

DNA metabarcoding is a powerful technique that can be harnessed to generate numerous sequence reads in parallel for multi-species identification and much more. Presently, DNA metabarcoding is conducted using second generation sequencing mainstays like Illumina, and less so on third-generation sequencers like ONT’s MinION sequencer. We surmised that this was likely due to the notoriously high error rates of nanopore reads, as well as the general lack of specialised programs that can process such erroneous reads. Existing nanopore metabarcoding workflows either incorporate complicated and time-consuming laboratory steps, or require custom reference databases, or additional polishing steps, which perhaps disincentives the use of nanopore sequencing for metabarcoding. However, recent improvements in nanopore read accuracy in conjunction with new bioinformatic pipelines have led us to posit that nanopore sequencing can now produce highly-accurate metabarcoding results that are consistent with conventional Illumina sequencing, and without the need to polish the sequences unlike in the past. We demonstrated this by metabarcoding 34 bulk zooplankton communities on two R10.3 MinION flow cells, and processed the reads with amplicon_sorter. Our results showed that: (1) nanopore metabarcodes are nearly Illumina-like in sequencing accuracy (99.6%) and are almost always indel-free (91.4%); (2) relative abundance of MOTUs were highly congruent (88%) across both platforms, and nanopore recovered the abundant MOTUs just as well as Illumina but struggled to capture the rarer taxa; and that (3) ecological conclusions were consistent across sequencing platforms when metabarcoding zooplankton communities despite some differences in species richness recovered. Reports of the newly released R10.4 sequencing chemistry already indicate vast improvements in the quality of nanopore sequences. We are confident that our results will inspire greater assurance in the utility of the MinION sequencer for more, and perhaps even larger-scale, metabarcoding-related projects in the near future.

## List of Abbreviations

MOTU: molecular operational taxonomic unit
NGS: next-generation sequencing
NMDS: nonmetric multidimensional scaling
ONT: Oxford Nanopore Technologies
PCR: polymerase chain reaction
PERMANOVA: permutational multivariate analysis of variance
SFB: short fragment buffer
SUP: super accurate
WoRMS: World Register of Marine Species

## Author Contributions

JJMC and DH conceived the project idea. MADM and ZJ led sample collections, assisted by WLN. WLN processed the samples and performed the wet laboratory processes together with YCAI. JJMC prepared the nanopore sequencing libraries, analysed the data and drafted the manuscript, with input from DH and YCAI. YCAI compiled the information for verification of taxonomic identities and geographic ranges. All authors reviewed the manuscript and approved the final draft for submission.

## Acknowledgements

We would like to thank Bing Jun Woo, Sarah Nelson, and Edwin Ong for their assistance with fieldwork and collections. We also acknowledge the National Supercomputing Centre (NSCC), Singapore and NUS High Performance Computing (HPC) for permitting the use of their computing resources for analyses, as well as the World Register of Marine Species (WoRMS) for making their data available to us.

## Data Availability

The Illumina sequence reads, nanopore base-called fast5 files, and nanopore fastq reads have been uploaded onto NCBI Sequence Read Archive under BioProject XXXXX. Sample metadata, demultiplexing information, OTU table, and taxonomic identifications can be found in Supplementary File S1.

## Figure Captions

**Figure S1.** Heatmap showing the pairwise Jaccard (top) or Bray-Curtis (bottom) distances between Illumina and nanopore samples. Values were converted to similarity (1 − dissimilarity score) for ease of understanding. Values close to 0 (blue) indicate no similarity, while values closer to 1 (red) indicate high similarity.

